# Gut Microbial Utilization of the Alternative Sweetener, D-Allulose, via AlsE

**DOI:** 10.1101/2024.11.07.622513

**Authors:** Glory Minabou Ndjite, Angela Jiang, Charlotte Ravel, Maggie Grant, Xiaofang Jiang, Brantley Hall

## Abstract

D-allulose, a rare sugar with emerging potential as a low-calorie sweetener, has garnered attention as an alternative to other commercially available alternative sweeteners, such as sugar alcohols, which often cause severe gastrointestinal discomfort. D-allulose-6-phosphate 3-epimerase (AlsE) is a prokaryotic enzyme that converts D-allulose-6-phosphate into D-fructose-6-phopshate, enabling its use as a carbon source. However, the taxonomic breadth of AlsE across gut bacteria remains poorly understood, hindering insights into the utilization of D-allulose by microbial communities. In this study, we provide experimental evidence showing that *Clostridium innocuum* is capable of D-allulose metabolism via a homologous AlsE. A bioinformatics search of 85,202 bacterial genomes identified 116 bacterial species with AlsE homologs, suggesting a limited distribution of AlsE in bacteria. Additionally, *Escherichia coli* contains a copy of *alsE*, but it does not grow on D-allulose as a sole carbon source unless *alsE* is heterologously expressed. A metagenomic analysis revealed that 15.8% of 3,079 adult healthy human metagenomic samples that we analyzed contained *alsE*, suggesting a limited prevalence of the enzyme in the gut microbiome. These results suggest that the gut microbiome has limited capacity to metabolize D-allulose via *alsE*, supporting its use as an alternative sweetener with minimal impact on microbial composition and gastrointestinal symptoms. This finding also enables personalized nutrition, allowing diabetic individuals to assess their gut microbiota for *alsE*, and manage glycemic response while reducing gastrointestinal distress.

## Introduction

The obesity epidemic is a serious health issue affecting many countries worldwide.^1^ According to the National Health and Nutrition Examination Survey (NHANES), conducted by the National Center for Health Statistics (NCHS), 41.9% of U.S. adults aged 20 and older are obese.^2^ As an individual’s amount of adipose tissue increases, so too does their risk for metabolic diseases, including type 2 diabetes,^3^ which is caused by insulin resistance and lack of insulin, resulting in chronic hyperglycemia.^4^ Over 415 million people worldwide suffer from diabetes, over 90% of whom have type 2 diabetes.^5^ In the U.S., 14.8% of adults aged 20 or older are also affected.^2^

Previous studies have linked increased sugar consumption to the obesity and diabetes epidemic.^6,7^ Further, researchers propose that a high-carbohydrate diet promotes the deposition of calories into fatty tissue, leading to weight gain through increased hunger.^8^ The main culprits of type 2 diabetes are excessive sugar consumption and a sedentary lifestyle.^5^ There is no cure for type 2 diabetes available as of 2024, and much more research is needed on methods to mitigate and prevent diabetes, including decreasing the consumption of fructose, glucose, and sucrose.

One potential strategy to minimize sugar consumption is to use sugar substitutes, such as aspartame, sucralose, erythritol, xylitol, and sorbitol.^9^ These alternative sweeteners tend to taste sweet, but the human body does not metabolize them, thereby reducing the adverse health effects of excess sugar consumption.^10^ These sweeteners may be derived from plant extracts or from chemical synthesis.^11^

Many of the alternative sweeteners currently approved by the U.S. Food and Drug Administration include sugar alcohols and non-nutritive sweeteners (NNS), both of which have been associated with some side effects on the human gut microbiome. Sugar alcohol consumption can lead to gastrointestinal discomfort and have laxative effects through osmotic pressure and increased gas production through gut bacterial fermentation, resulting in diarrhea and bloating.^12^ Moreover, increased blood erythritol levels have been associated with increased platelet reactivity, resulting in cardiovascular events such as strokes.^13^ On the other hand, regular NNS consumption can lead to functional alteration of gut microbiota composition, resulting in an impaired glycemic response and glucose intolerance.^14,15^ Therefore, it is extremely important to understand the mechanisms of how alternative sweeteners interact with the human gut microbiome. These findings have led to an increasing interest in fructose epimers, sugar molecules that resemble fructose but have altered stereochemistry at one carbon atom.^16^ One example is D-allulose (also known as D-psicose), a rare low-calorie sweetener that is the C-3 epimer of fructose and is found in small amounts in certain fruits.^16^ Previous studies have suggested that D-allulose has a low glycemic index, making it promising for reducing the risk of diabetes.^17–20^ Due to advances in the industrial process and bacterial engineering methods, D-allulose production is becoming increasingly economically viable.^21^ Thus, D-allulose is a promising way to decrease sucrose and fructose consumption.

Although D-allulose is a promising alternative sweetener, its side effects are poorly understood compared to other types of alternative sweeteners. A significant portion of ingested D-allulose reaches the gut microbiome, as approximately 30% passes through the small intestine unabsorbed and is excreted in feces^22,23^. While 70% of D-allulose is absorbed via glucose transporter type 5 (GLUT5) in the small intestine, the substantial unabsorbed fraction has the potential to interact with and impact the gut microbial community.^22,23^ Studies in murine models have shown that D-allulose can induce changes in the gut microbiome.^24,25^ Comparatively, Suez et al. 2014 showed that saccharin, sucralose, and aspartame can induce glucose intolerance through modifications of the gut microbiome composition and function.^26^ Therefore, there is an urgent need to better understand both the potential for D-allulose utilization by gut bacteria and its effects on human gut microbiome composition.

Some bacteria possess the ability to metabolize D-allulose using the enzyme D-allulose-6-phosphate 3-epimerase (AlsE), encoded by the gene *alsE*.^27^ In *E. coli* K-12, *alsE* is in the D-allose operon, which has been well characterized (**Figure 1A**).^28^ First, D-allose is converted into D-allulose 6-phosphate via AlsK and RpiB. Then, AlsE catalyzes the reversible conversion of D-allulose 6-phosphate to D-fructose 6-phosphate.^29^ Environmental and clinical isolates of *Klebsiella pneumoniae,* an opportunistic pathogen responsible for a significant number of nosocomial bacterial infections,^30^ are capable of metabolizing D-allulose using a homologous AlsE, raising concerns that consuming D-allulose may confer opportunistic pathogens an advantage in colonization.^31,32^ However, there has not been a full systematic annotation of the prevalence and distribution of *alsE* in the human gut microbiome.

**Figure 1:**
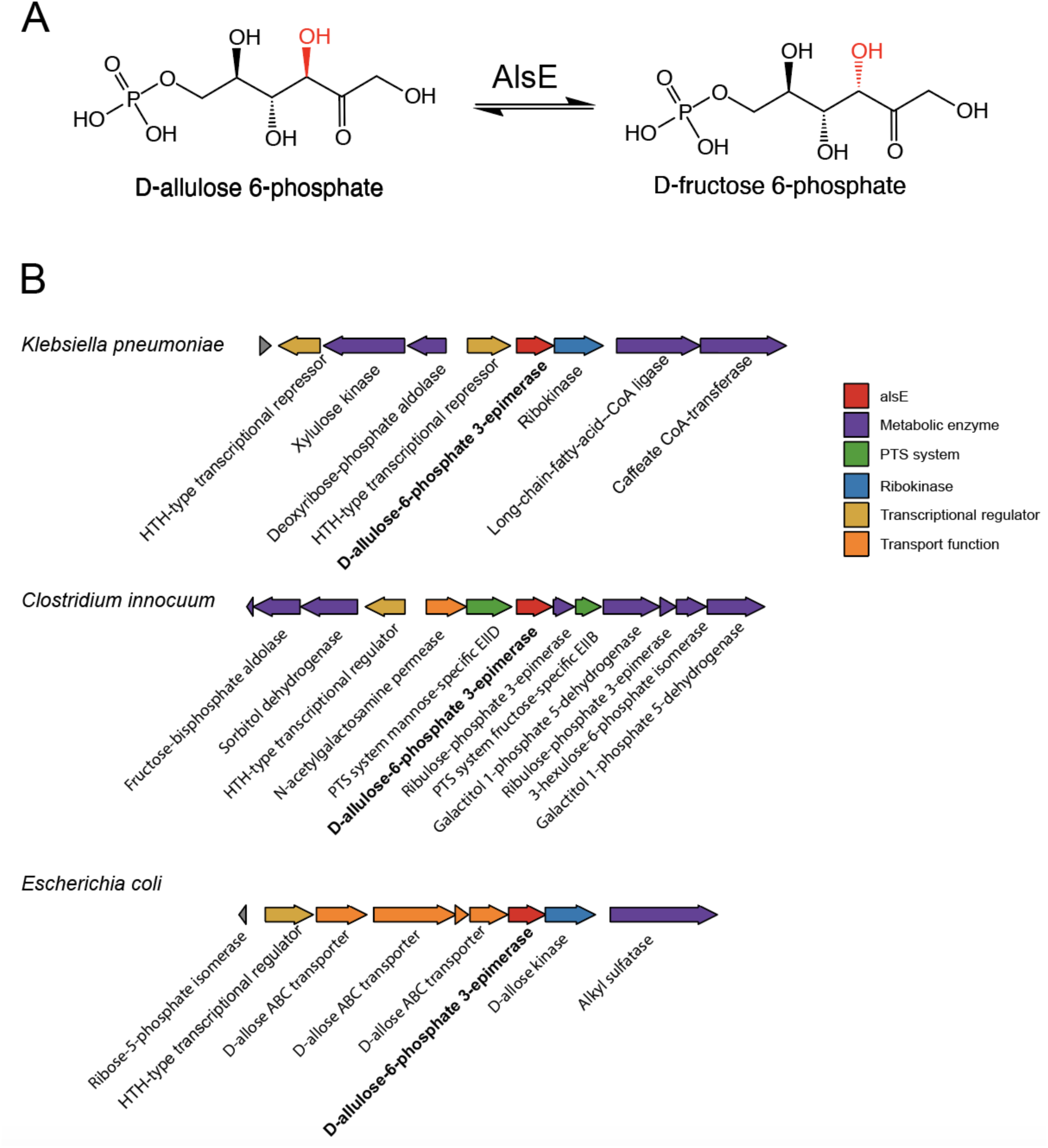
A) Conversion of D-allulose-6-phosphate into D-fructose-6-phosphate by *alsE*. B) *alsE* cluster organization from *Klebsiella pneumoniae* MGH78578, *Clostridium innocuum* 6_1_30, and *Escherichia coli* K-12.

To address this gap, we combined bioinformatic predictions and experimental verification to characterize the distribution of *alsE* in human gut microbes. Our bioinformatic predictions were validated through growth experiments, culturing bacteria in media with D-allulose as the sole carbon source. Our investigation of multiple representatives of the major gut bacterial clades expanded the known phylogenetic range of D-allulose metabolism from phylum Pseudomonadota to include phylum Bacillota by identifying that *Clostridium innocuum* 6_1_30 is capable of using D-allulose as a sole carbon source. Through comparative genomics and protein homology searches, we identified a putative AlsE in *C. innocuum* that is homologous to AlsE in *K. pneumoniae* (35% identity, e-value 2.89e-41) that has a divergent operon organization compared to the known D-allulose metabolizers. We verified the function of the *C. innocuum* AlsE in the *E. coli* Keio Knockout Collection, observing that *E. coli* deficient in native *alsE* was able to grow on D-allulose when complemented with *C. innocuum alsE*. We also found that *E. coli*, despite encoding *alsE*, cannot grow on D-allulose as a sole carbon source unless *alsE* is heterologously expressed. To comprehensively characterize the taxonomic distribution of AlsE, we performed a systematic search across 85,202 bacterial genomes, identifying 116 species encoding putative *alsE* homologs. The limited distribution of *alsE* in the gut microbiome supports D-allulose’s promise as an alternative sweetener with minimal impact on both microbial composition and gastrointestinal symptoms, two common drawbacks of current artificial sweeteners. Although our focus is on *alsE*, it is important to note that there could be alternative undiscovered pathways bacteria can use to metabolize D-allulose that our study did not cover. These findings provide insights into bacterial D-allulose metabolism, supporting its development as an alternative sweetener to help reduce sugar consumption in the context of rising rates of obesity and diabetes.

## Results

### Investigating the potential for D-allulose utilization by gut microbes

To identify gut bacterial species capable of utilizing D-allulose as a carbon source, we conducted a preliminary identification of species with D-allulose-6-phosphate 3-epimerase (AlsE) homologs by conducting a BLASTp search against 85,202 non-redundant genomes from the Genome Taxonomy Database^33,34^. We used the experimentally verified *Klebsiella pneumoniae* AlsE as the query, with a threshold of 50% identity and bitscore greater than 200. There were 272 species that met the threshold, mostly from non-gut bacteria. Some gut bacteria genera with AlsE homologs include *Klebsiella*, *Escherichia*, and *Clostridium*. Interestingly, *Clostridium innocuum*, a common gut bacterium species, contained an AlsE hit (50.49% identity, 2.30e-77 e-value, 220 bitscore).

To experimentally validate our bioinformatic predictions and characterize D-allulose metabolism across diverse gut bacteria, we tested representative strains from major gut bacterial phyla including Bacillota, Bacteroidota, Actinobacteriota, and Pseudomonadota for growth on D-allulose as a sole carbon source (**Figure 2A**). Growth was quantified by spectrophotometric measurement at OD600, with significant growth defined as a three-fold increase in OD600 compared to media negative controls. *Clostridium innocuum* 6_1_30 demonstrated robust growth on D-allulose with a 6:1 ratio in its OD600 measurement compared to the media blank, revealing a previously unknown metabolic capability (**Figure 2B**). Notably, *Escherichia coli* DC10B did not grow on D-allulose despite encoding *alsE* within its D-allose operon (**Figure 2C**), prompting further investigation.

**Figure 2:**
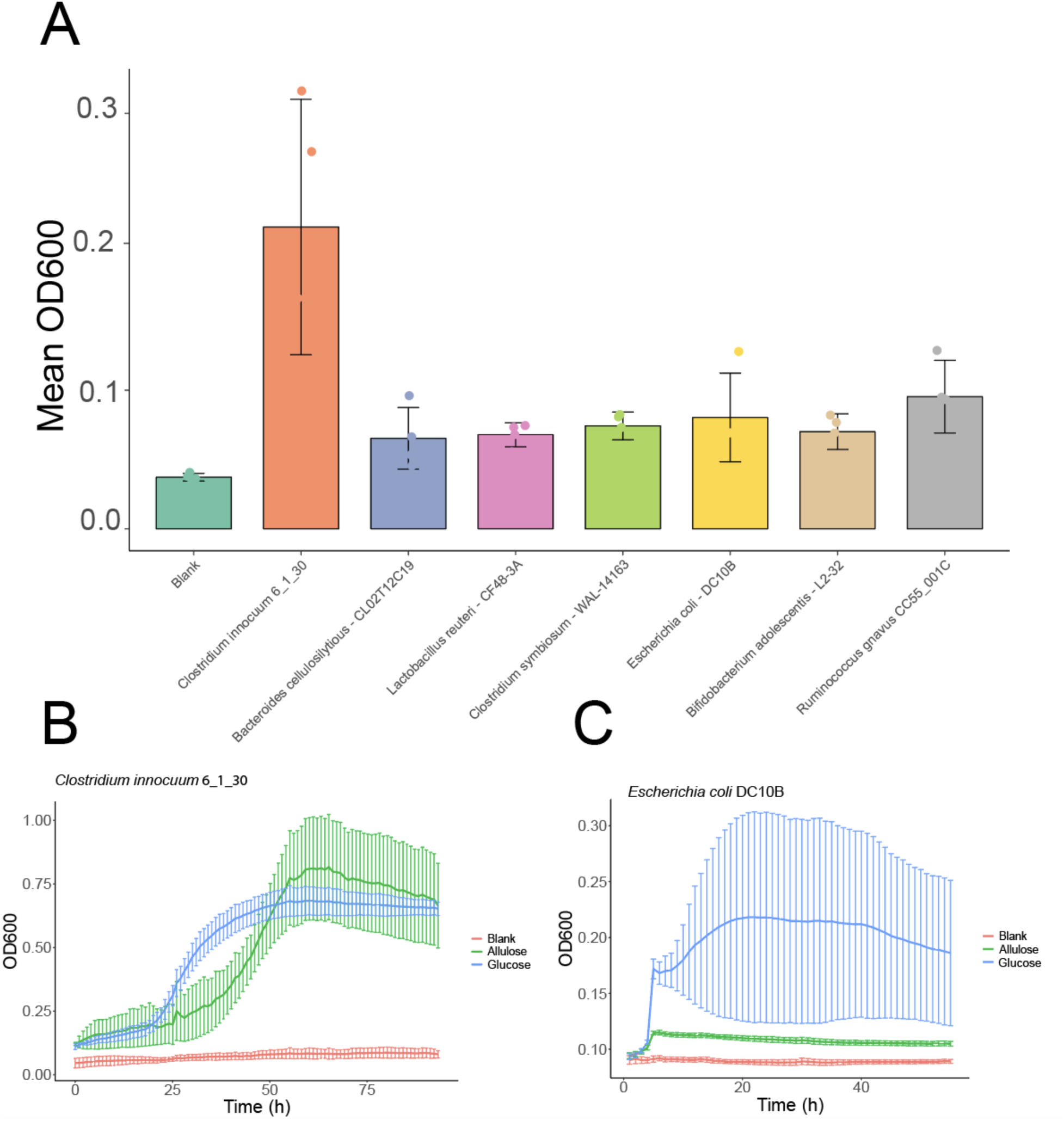
Identification of *Clostridium innocuum* 6_1_30 as a gut bacteria species that can grow on allulose as a sole carbon source. A) Investigation of 7 gut bacteria species (*Clostridium innocuum* 6_1_30, *Bacteroides cellulosilytious* - CL02T12C19, *Lactobacillus reuteri* CF48-3A, *Clostridium symbiosum* WAL-14163, *Escherichia coli* DC10B, *Bifidobacterium adolescentis* L2-32, *Ruminococcus gnavus* CC55_001C) for growth on allulose as the sole carbon source. Each data point is the average of three technical replicates from a single biological replicate per species. B) Growth curve of *C. innocuum* 6_1_30 on allulose with minimal media. Allulose is the growth curve of *C. innocuum* when grown on allulose, while Glucose is a positive control of *C. innocuum* growing on glucose, and Blank refers to *C. innocuum* grown on blank media as a negative control. C) Growth curve of *E. coli* DC10B (Col02) on allulose with minimal media.

### *Escherichia coli* does not readily utilize D-allulose *in vitro* despite encoding *alsE*

Although *E. coli* encodes *alsE* within the D-allose operon (*alsRBACEK*), previous studies have demonstrated that this operon is specifically induced in response to D-allose.^28^ We hypothesized that while *E. coli* encodes the metabolic machinery for D-allulose utilization; this capability may not be active in the absence of D-allose.

To test this hypothesis, we placed *alsE* under the control of an IPTG-inducible promoter to enable controlled expression independent of its native regulation. Using the Keio collection, a comprehensive library of single-gene knockout mutants in *E. coli* BW25113^35^, we cloned the *alsE* gene from strain JW2760 into a pCW-lic vector backbone under an inducible *tac* promoter, creating the pCW-lic-*E.coli alsE* construct. This plasmid was transformed into the Keio *alsE* knockout strain, and the transformed bacteria were cultured in M9 minimal media supplemented with D-allulose and IPTG.^12^

Our results showed that both the Keio *alsE* knockout and the untransformed *E. coli* were unable to grow using D-allulose as the sole carbon source. In contrast, the transformed *E. coli* overexpressing *alsE* exhibited robust growth (**Figure 3A and 3B**). These findings support our hypothesis that while *E. coli* encodes AlsE, it may not be capable of utilizing D-allulose as a carbon source in the absence of D-allose.

**Figure 3:**
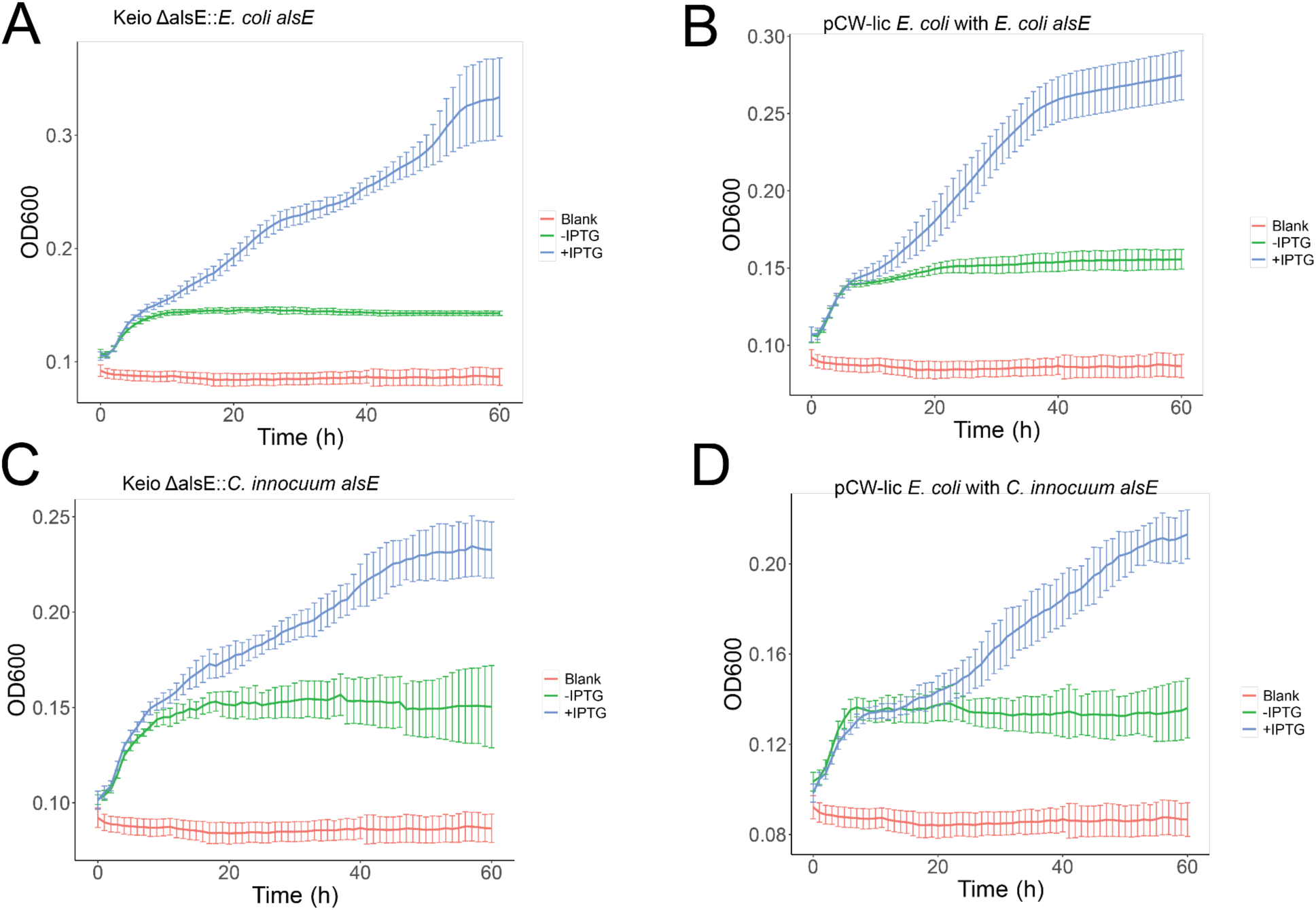
A and B) Verification of the *Escherichia coli alsE* functionality. Keio Δ*alsE*::*E. coli alsE* + IPTG is the growth curve of the transformed Keio pCW-lic-E.coli_*alsE*, containing the wild-type *E. coli alsE* with IPTG to induce ectopic expression. Keio Δ*alsE*::*E. coli alsE* is the growth curve of the transformed Keio *E. coli* without IPTG, resulting in no gene expression. C and D) Verification of the *Clostridium Innocuum* 6_1_30 *alsE* functionality. Keio Δ*alsE*::*C. Innocuum alsE* + IPTG is the growth curve of the transformed Keio pCW-lic_C.Inn_*alsE*, containing the *C. innocuum alsE* with IPTG to induce ectopic expression. Keio Δ*alsE*::*C. Innocuum alsE* is the growth curve of the same transformed Keio *E. coli* without IPTG, resulting in reduced gene expression.

### Limited distribution of *alsE* across *E. coli* strains

We then investigated the presence and absence of *alsE* across *E. coli* genomes, using a previously published pangenome consisting of 1,324 *E. coli* genomes.^36^ *alsE* was present in 598 out of 1324 *E. coli* genomes (45%), suggesting that *alsE* is a strain-specific gene and may not be present in every individual’s gut microbiome, despite the prevalence of *E. coli* exceeding 90% among in humans.^37^

### Identification of alsE in Clostridium innocuum

Given that *Clostridium innocuum* 6_1_30 grew on D-allulose as a sole carbon source, we investigated the genomic origins of its D-allulose metabolism. Based on our prior preliminary search results, we hypothesized that a homologous *alsE* was responsible for D-allulose metabolism in *C. innocuum* rather than a novel pathway. To conduct a comprehensive search for AlsE homologs in *C. innocuum*, we used the *Klebsiella pneumoniae* MGH 78578 AlsE (NCBI accession: GCA_000016305.1) as the query to search the *C. innocuum* genome (NCBI accession: GCA_000183585.2). BLASTp revealed two AlsE homologs in *C. innocuum*, referred to as ci04257 and ci04568 (ci04257: 50.49% identity, 2.09e-76 e-value; ci04568: 35% identity, 2.89e-41 e-value). We examined the gene neighborhood of the two *alsE* candidates. The neighborhood of ci04257 consisted mainly of genes encoding hypothetical proteins. On the other hand, ci04568 was adjacent to phosphotransferase systems (PTS), which could potentially perform the phosphorylation and import step of D-allulose utilization. In addition, the neighboring genes are annotated with sugar metabolism functions, such as fructose bisphosphate aldolase. Therefore, we hypothesized that ci04568 encodes an enzyme that possibly performs a similar function to AlsE. Interestingly, the putative *alsE* gene neighborhood in *C. innocuum* is completely divergent from the *alsE* gene neighborhood in other species known to metabolize D-allulose, such as *Klebsiella pneumoniae* (**Figure 1B**).^31^ Of note, this putative *alsE* was a core gene present in all 283/283 available *C. innocuum* genomes on NCBI, with all of them containing a nearly identical, if not identical, homolog to *alsE* in 6_1_30.

We then sought to functionally validate the candidate *alsE* in *C. innocuum* by cloning the gene into a pCW-lic vector backbone under an inducible tac promoter, resulting in the pCW-lic_C.inn_*alsE* construct to heterologously express *C. innocuum’s alsE* in *E. coli* (**Supplemental Figure 1**). The plasmid was then transformed into the Keio collection *E. coli alsE* knockout. The transformed bacteria were subsequently inoculated in D-allulose-supplemented M9 and induced *alsE*’s expression using IPTG.^12^ The Keio *alsE* knockout demonstrated no growth on D-allulose, while complementation of the *C. innocuum* gene into the knockout restored function, resulting in growth on D-allulose (**Figure 3C and 3D**).

### Few Gut Bacterial Species Encode AlsE

Once we experimentally verified the function of the *C. innocuum* and *E. coli* AlsE, we used ProkFunFind,^38^ a bioinformatics pipeline to systematically search for AlsE in bacteria. We used the experimentally verified AlsE protein sequences from *K. pneumoniae* MGH 78578, *C. innocuum* 6_1_30, and *E. coli* K-12 as queries to search the 85,202 non-redundant prokaryotic genomes from the Genome Taxonomy Database (GTDB) for species that contained homologs to AlsE. We used a more stringent filtering criteria compared to the preliminary search, filtering hits based on a 30% identity threshold and a maximum e-value of 1e-100. Our search revealed 116 putative bacterial species with AlsE (**Supplemental Table 3**, **Figure 4**). The vast majority of these species were from the phylum Pseudomonadota (103/116), 10 were from the phylum Bacillota, and 3 were from Fusobacteriota. Out of those 116 species, only 35 are known to be part of the animal gut microbiota. Some known members of the human gut microbiome with AlsE include *Klebsiella oxytoca*, *Enterobacter cloacae*, and *Serratia marcescens*. Other species with AlsE that are not gut-associated are primarily isolated from plants and soil, including *Klebsiella planticola*,^39^ *Rahnella aquatilis*,^40^ and members of the *Kosakonia* genus.^41,42^ Of note, none of the species that were unable to grow on D-allulose during our initial investigation contained AlsE homologs, with the exception of *Escherichia coli* as discussed before (*Bacteroides cellulosilytious*, *Lactobacillus reuteri*, *Clostridium symbiosum*, *Bifidobacterium adolescentis*, *Ruminococcus gnavus*).

**Figure 4:**
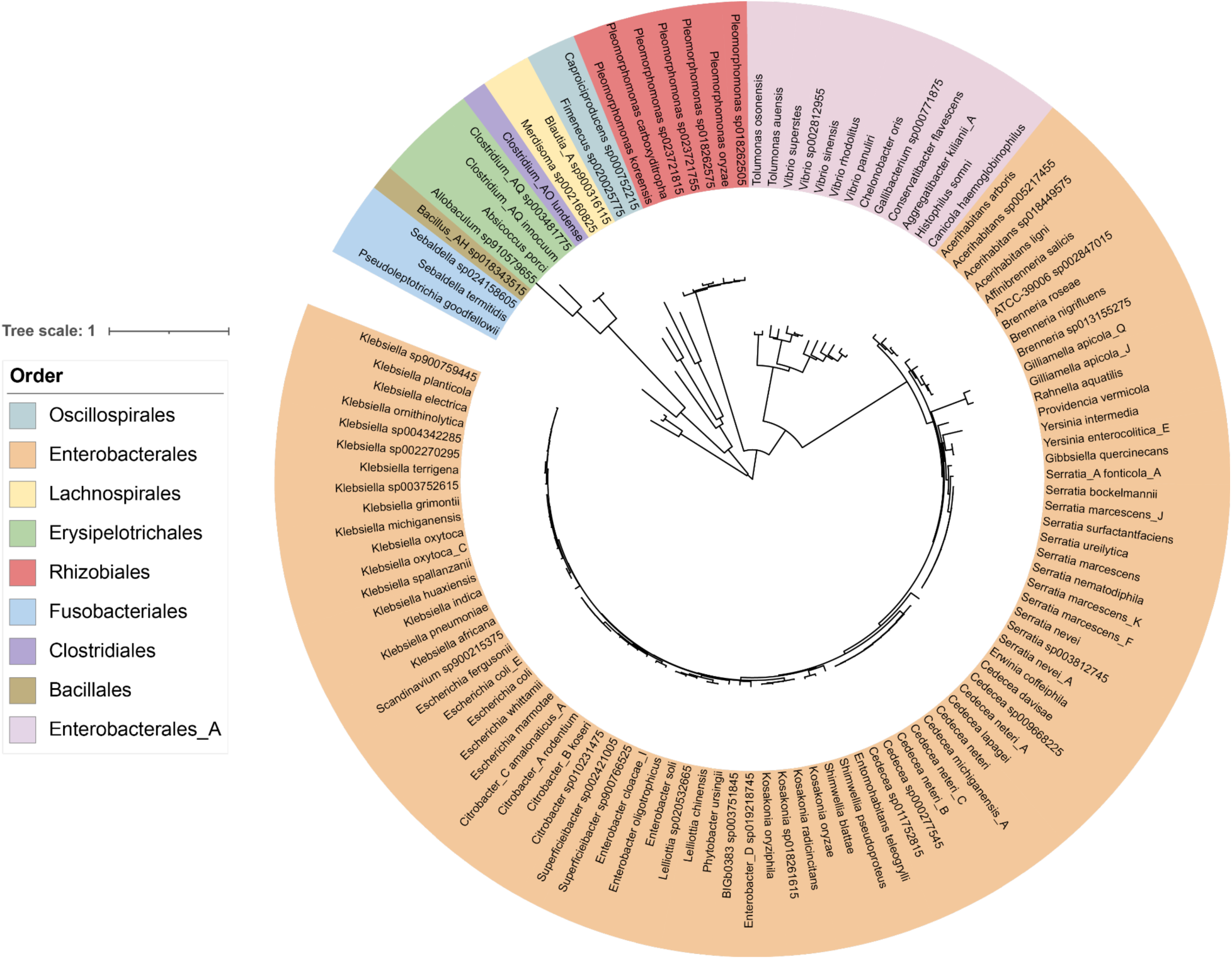
Species tree showing the taxonomic distribution of AlsE in microbial genomes from GTDB, colored by order. The species tree was generated by pruning the GTDB species tree using the Gotree prune command.^51^

### Presence of *alsE* in the healthy adult gut microbiome

To investigate the prevalence of *alsE* in the human gut microbiome, we examined the presence and absence of *alsE* in 3,079 healthy adult human gut microbiomes that passed quality checks. We built a reference database using both experimentally characterized and bioinformatically discovered *alsE*, with thresholds of 30% identity and 1e-100 e-value. We then aligned healthy adult human stool metagenomic reads downloaded from SRA to our *alsE* reference database and normalized the alignment counts into counts per million. To strike a balance between spurious hits and sensitivity, we considered any metagenomes with at least 1 count per million to contain *alsE*. 488 out of 3,079 metagenomes met our threshold for *alsE* presence, approximately 15.8%.

### Delineation of AlsE from Pentose-5-Phosphate 3-Epimerase

In order to elucidate the evolutionary origins of AlsE, we used a combination of phylogenetic analyses, ancestral state reconstruction, and sequence conservation. Using eggNOG-mapper (v6.0), we determined that AlsE belonged to the orthologous group COG0036 (pentose-5-phosphate 3-epimerase). We constructed a phylogenetic tree using the top 1,323 homologs of the 3 experimentally verified AlsE protein sequences against all COG0036 sequences. Based on branch lengths and the presence of the experimentally verified AlsE, we identified a putative AlsE clade that contains 515 sequences. We were able to identify conserved amino acid changes in the putative AlsE node from the ancestral node (**Figure 5A**), such as from G52 to S52, V134 to Y134, L142 to T142, and an N147 to D147 **(Figure 5B)**. Based on the high entropy in the alignment at these positions, these are conserved changes and may differentiate AlsE from other pentose-5-phosphate 3-epimerases.

**Figure 5:**
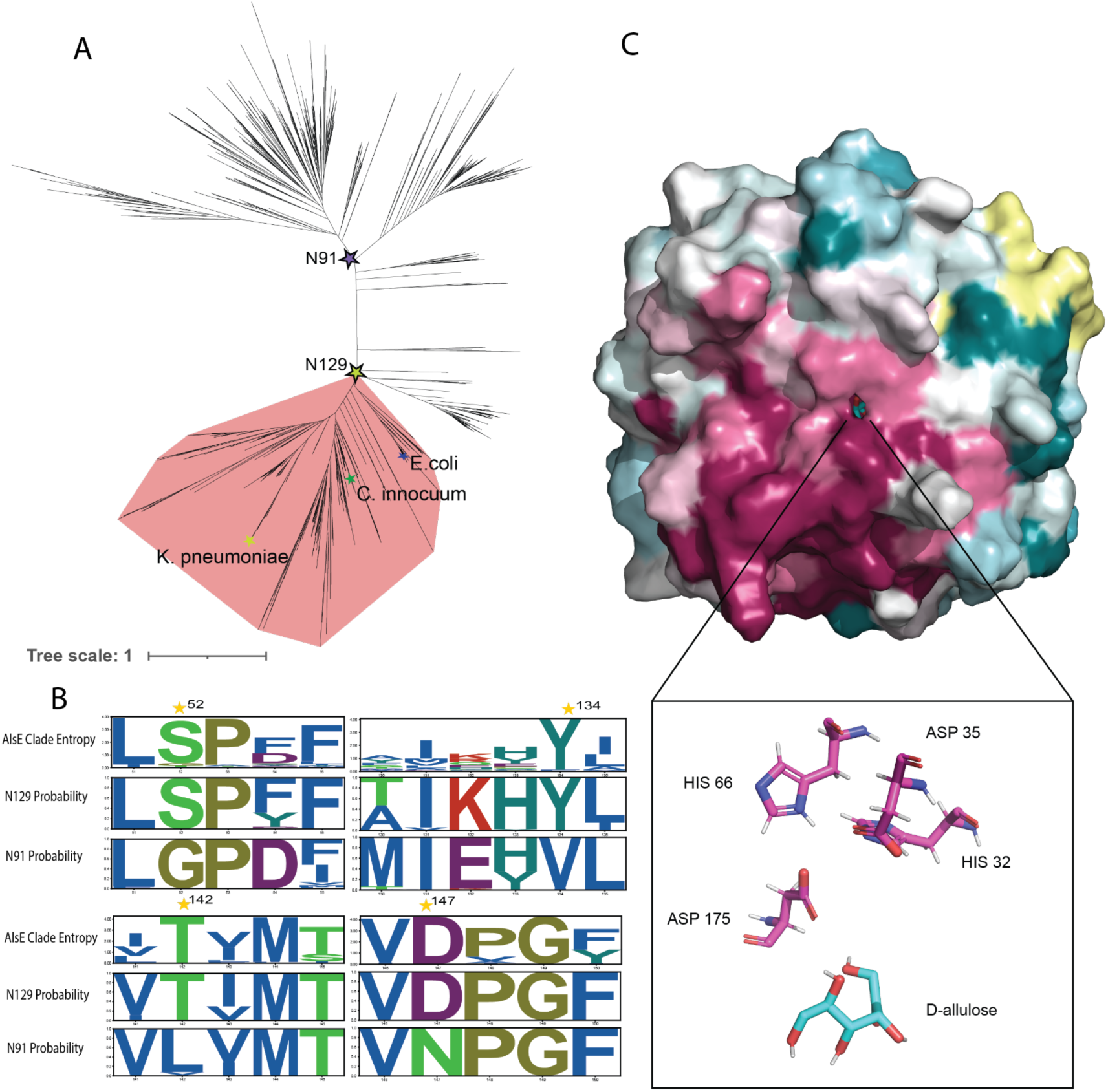
A) Gene tree constructed from putative AlsE sequences and related enzymes annotated as pentose-5-phosphate-3-epimerase (COG0036), showing a possible delineation of the AlsE clade and the location of *Klebsiella pneumoniae*, *Clostridium innocuum*, and *Escherichia coli* AlsE. B) Diagram showing the change and conservation (entropy) of residues in the putative AlsE clade, as well as the GRASP predicted ancestral states of N129 (AlsE clade) and N91 (putative ancestral node). Residues with a predicted conserved change from the ancestral state are labeled with a star. C) D-allulose docked to the AlphaFold2-predicted structure of *C. innocuum* 6_1_30 AlsE, colored by amino acid conservation via Consurf.

To determine putative catalytic residues, we performed a structural alignment of the AlphaFold2-predicted structure of *C. innocuum* AlsE to the crystal structure of *E. coli* AlsE (pdb: 3CT7). Based on the active site residues reported by Chan et al. 2008 in *E. coli* AlsE, we determined that the putative active site residues in both *Clostridium innocuum* and *Klebsiella pneumoniae* are similar based on the structural alignment (**Figure 5C**).^29^ In *C. innocuum*, these putative residues are His 32, Asp 35, His 66, and Asp 175, which align to His 34, Asp 36, His 67, and Asp 176 in *E. coli*, respectively. Therefore, despite being distant homologs, *C. innocuum* likely shares similar catalytic residues to *E. coli* AlsE.

## Discussion

Many widely used commercial alternative sweeteners, such as sugar alcohols, are associated with significant gastrointestinal discomfort.^12^ This discomfort arises from the malabsorption of these sweeteners, leading to osmotic diarrhea, and from fermentation by gut microbes, which produce gas.^12^ Consequently, there is an urgent need to identify alternative sweeteners, such as D-allulose, that do not cause gastrointestinal symptoms.

The presence of gut bacteria that can potentially metabolize D-allulose via D-allulose-6-phosphate 3-epimerase (AlsE) has significant implications for its use as a commercial alternative sweetener. Prior to our study, while D-allulose metabolism had been identified in some human gut bacteria, there had not been a systematic analysis of the presence, abundance, and distribution of enzymes involved in D-allulose metabolism across gut bacterial species - a knowledge gap that limited our understanding of how gut bacteria utilize this sweetener. In our study, we demonstrated that *Clostridium innocuum* can metabolize D-allulose through a homologous AlsE by examining its growth on D-allulose media. These findings shed light on the role of the gut microbiome in D-allulose metabolism.

During the initial investigation for gut microbial species capable of growing on D-allulose as a sole carbon source, *C. innocuum* 6_1_30 grew on D-allulose as a sole carbon source while *E. coli* was unable to grow despite encoding *alsE* in its genome, which was intriguing. Past studies have shown that despite *E. coli* having *alsE* in its genome, its expression was too weak to support the production of D-allulose from D-fructose without genetic modifications.^21,43^ This is consistent with our findings that the *E. coli* only grew on D-allulose when *alsE* was heterologously expressed, verified via the insertion of the respective *alsE* genes into the Keio *alsE* knockout mutant, which resulted in *E. coli* gaining the ability to use D-allulose as a sole carbon source.

Our findings show that AlsE protein homologs are only present in a few gut bacterial species. Out of 85,202 bacterial genomes from the GTDB, only 116 bacterial species were annotated to contain AlsE homologs. Our finding that *E. coli* cannot grow on D-allulose without heterologously expressing *alsE* suggests that some of these 116 species may not be able to metabolize D-allulose effectively. In addition, only 35 of these species are known to be present in animal gut microbiomes. These data suggest that D-allulose utilization might be restricted to a small number of species within the human gut microbiome. This finding is in alignment with the scarcity of D-allulose in nature. D-allulose has only been found in small quantities in a few plant species, such as *Itea virginica* and wheat.^44,45^ Moreover, several of the bacterial species with putative *alsE* were primarily isolated from plants such as wheat or maize, including *Klebsiella planticola*,^39^ *Rahnella aquatilis*,^40^ and members of the *Kosakonia* genus.^41,42^ We speculate that D-allulose metabolism may confer a metabolic advantage for these bacteria that live in plant-associated habitats, where exposure to D-allulose is more likely. Alternatively, *alsE* may have evolved primarily to confer D-allose metabolism, with D-allulose metabolism being incidental.

Notably, the limited presence of *alsE* in gut microbiome species suggests that D-allulose may serve as a valuable alternative to common sugar substitutes, which are known to cause gastrointestinal discomfort and alter microbiome composition. Previous studies have reported that D-allulose can be consumed in relatively high doses, up to 0.5 g/kg body weight, without causing significant gastrointestinal issues.^46^ Thus, the limited metabolism of D-allulose by gut bacteria, combined with its low impact on gastrointestinal function, suggests that it may offer a promising solution for individuals seeking low-calorie sweeteners without adverse digestive effects. Of note, approximately 15.8% of human metagenomes analyzed contained *alsE*, suggesting that individual gut microbiomes may respond differently to D-allulose consumption. Diabetic individuals looking to cut their glucose consumption may benefit from individual microbiome testing to choose the alternative sweetener that is less likely to be utilized by their gut microbiome.

Our study focuses on AlsE as an enzyme responsible for D-allulose metabolism, though we recognize the possibility of alternative mechanisms of D-allulose metabolism. To our knowledge, AlsE is the only currently known enzyme implicated in D-allulose metabolism in bacteria. However, there may be alternative mechanisms of bacterial D-allulose metabolism that are undiscovered, given the limited studies on the subject.^28,31^ Due to this possibility of unknown alternative mechanisms, we cannot be certain of D-allulose’s impact on gut microbiome composition at large.

In conclusion, we shed light on the taxonomic distribution of AlsE in the gut microbiota. We discovered that *C. innocuum* is capable of growing on D-allulose as a sole carbon source. In addition, while *E. coli* has *alsE*, it cannot grow on D-allulose without heterologously expressing *alsE*, suggesting that many of these bacteria do not necessarily grow on D-allulose as a sole carbon source. A relatively small fraction of gut microbes are capable of utilizing D-allulose, making it a promising alternative to commercially available sugar substitutes, such as sugar alcohols.

## Methods

### Identification of D-allulose-6-phosphate 3-epimerases in the GTDB genomes

All representative genomes from the Genome Taxonomy Database (GTDB) (release r207) were downloaded, and protein sequences for each genome were predicted using Prokka (version 1.14.6).^34,47^ The *Escherichia coli* K-12, *Klebsiella pneumoniae* MGH78578, and *Clostridium innocuum* 6_1_30 D-allulose-6-phosphate 3-epimerase protein sequences were searched against 85,202 reference genomes using the ProkFunFind pipeline (v0.1.0).^38^ The hits were filtered based on an 1e-100 e-value and 30% identity thresholds, resulting in putative 126 AlsE amino acid sequences from 116 nonredundant genomes.

### Phylogenetic analyses

Sequences from the GTDB assigned to COG0036 were identified using eggNOG-mapper (version 2.1.3).^48^ A BLASTp search was conducted (version 2.15.0+) using the identified D-allulose 6-phosphate 3-epimerases as queries against these identified sequences, setting a limit to the top 1,305 hits. Sequence alignment was performed using Clustal Omega (version 1.2.4).^49,50^ Columns that have more than 97% gaps were removed to enhance alignment quality using Goalign (version 0.3.7).^51^ Phylogenetic analysis was carried out using IQ-TREE (version 2.1.2) with default parameters and model selection^52^. The reliability of the phylogenetic trees was evaluated using 1,000 ultrafast bootstrap replicates. Trees were visualized using the Interactive Tree Of Life (iTOL).^53^

Ancestral sequence reconstruction was performed on the AlsE tree using GRASP (version 04-May-2023), with default parameters.^54^ We then manually inspected the tree to delineate AlsE from other Pentose-5-phosphate 3-epimerases. We calculated the entropy of the alignments using Goalign via the compute pssm function (v.0.3.7).^51^ The figures were created using the Python package logomaker (v0.08).^55^

### Growth of anaerobic bacteria

Bacterial strains were acquired from the NIH Biodefense and Emerging Infections Research Resources Repository (BEI). Each strain was inoculated from a glycerol stock and grown under anaerobic conditions over a 24-hour period at 37 °C in an anaerobic chamber (Coy Laboratory Products) in Brain-Heart Infusion (BHI) broth (Research Products International, B11000) supplemented with glucose. 25 µL of the culture was inoculated into 4 mL of minimal media (M9) supplemented with 10 mg/mL carbon source (glucose or D-allulose).^12^

*Absorbance Assay:* The transformed Keio Δ*alsE*::*C. Innocuum alsE* & Keio Δ*alsE*::*E. coli alsE* constructs were shaken in Luria-Bertani (LB) supplemented with 100 µg/mL carbenicillin (GoldBio, C-103-25) overnight at 37°C. 25 µL of the overnight culture was inoculated in 4 mL triplicates of minimal media (M9) supplemented with 100µM Isopropyl β-d-1-thiogalactopyranoside (IPTG, GoldBio, I2481C25), 100 µg/mL carbenicillin, 50 µg/mL kanamycin (Bio Basic, KB0286), and 10 mg/mL D-allulose^12^ (Chem-Impex, 32353). For kinetic measurements, 250 µL of the triplicates were aliquoted into a 96-well acrylic, clear bottom plate (Celltreat, 229592), sealed with a Breathe Easy membrane (Electron Microscopy Sciences, 70536-10), and incubated at 37°C for 48-70 hours depending on the strain observed. The end-point absorbance at 600 nm was measured with a Spectramax M5 plate reader, with end-point bacterial growth calculated using a ratio to the blank, with a ratio of 3 indicating significant growth.

### pCW-lic_C.inn_alsE & pCW-lic-E.coli_alsE constructs

In order to achieve ectopic expression of *alsE* from *Clostridium innocuum* 6_1_30 and *E. coli* JW2760^35^ in the knockout mutants, the *alsE* gene was amplified and cloned into the pCW-lic vector backbone (Addgene, 26098). Genomic DNA from *C. innocuum* and *E. coli* JW2760 was utilized in a polymerase chain reaction (PCR) using Phusion High-Fidelity DNA Polymerase (NEB, M0530S) with the specific primers listed in Supplementary Table 1. A Monarch PCR & DNA Cleanup Kit (NEB, T1030S) was used to purify the amplified product. The pCW-lic vector backbone was digested with restriction enzymes NdeI (NEB, R0111S) and HindIII-HF (NEB, R3104S), followed by purification with a Monarch PCR & DNA Cleanup Kit. A Gibson assembly was completed using Gibson Assembly Master Mix (NEB, E2611S) in accordance with the manufacturer’s instructions. The resulting constructs were stored at -20°C until needed for use.

### Keio-pCW construct

The *alsE* gene was amplified and cloned into the pCW-lic vector backbone under a tac promoter and transformed into the Keio collection *als*E knockout as detailed above with the same primers outlined in Supplementary Table 1. For the control, an empty pCW-lic vector was cloned into the Keio *alsE* knockout.

### Chemical competency

The Keio collection *alsE* knockout was made competent using the Mix & Go! *E. coli* Transformation Kit and Buffer Set (Zymo, T3001) in accordance with the manufacturer’s protocol and stored at -80°C until needed for use.

### Transformation

Both constructs were independently transformed into the chemically competent Keio collection *alsE* knockout in accordance with the manufacturer’s protocol (Zymo, T3001). The resulting transformed cells were plated on LB agar plates supplemented with 100 µg/uL of carbenicillin. Successful transformation was validated via Oxford Nanopore sequencing by Plasmidsaurus.

### Structural prediction and molecular docking

The structure for the *Clostridium innocuum* 6_1_30 AlsE was predicted using AlphaFold2 (v2.3.0).^56^ Binding pockets were predicted using fpocket (v4.0) with default parameters.^57^ The pockets were compared to the homologous *Escherichia coli* AlsE (3CT7) to identify putative substrate binding regions and catalytic residues.^29^ The structure for D-allulose (PubChem compound identifier: 50909805) was docked onto the predicted AlsE structure using AutoDock Vina (v4.2).^58,59^ The docking simulation was performed within 15 Å × 15 Å × 15 Å cubes centered on the center points of the chosen fpocket substrate binding pocket with exhaustiveness set to 32. Docking results were visualized using PyMOL.^60^ We used Foldseek to identify the top structural homolog.^61^ The predicted AlsE protein structure was aligned with the *E. coli* 3CT7, and the putative catalytic residues were identified based on the previous work by Chan et al. 2008, using TM-Align.^29,62^ Protein sequence conservation of AlsE was visualized using ConSurf based on the putative AlsE clade.^63,64^

### Profiling of alsE presence in the gut

To build the reference database, we used *alsE* identified by ProkFunFind, which were filtered based on a threshold of e-value 1e-100 and percent identity 30%, resulting in a total of 126 sequences. We downloaded a collection of adult healthy metagenomic biosamples that passed basic quality control (n=3410) from SRA, and then trimmed adapters with Trim-Galore with default settings. The reads were then mapped to a human reference (assembly T2T-CHM13v2.0) to identify potential contaminants and removed them using Samtools (v1.16).^65^ We removed any samples with less than a million reads after curation, resulting in 3,079 samples, and then aligned the remaining reads to the *alsE* reference database using bowtie2 (v2.4.1).^66^ The number of reads mapped to the *alsE* reference was summarized by normalizing the number of reads in the sample and then multiplying by one million to obtain counts per million (cpm). If a biosample had multiple SRRs, we concatenated the read counts and total reads across all SRRs per sample before calculating cpm. We considered samples with at least 1 cpm as containing *alsE*, to account for spurious alignments.

## Author Statements

### Author contributions

B.H. and X.J. conceptualized and supervised the project. All authors performed the experiments and analyzed the data. G.M.N, A.J., C.R., and M.G. wrote the original draft of the manuscript. All authors reviewed and edited the paper.

## Acknowledgments

This work utilized the computational resources of the NIH HPC Biowulf cluster (http://hpc.nih.gov) and the UMIACS cluster at the University of Maryland’s Center for Bioinformatics and Computational Biology (https://www.umiacs.umd.edu/). pCW-LIC was a gift from Cheryl Arrowsmith (Addgene plasmid # 26098; http://n2t.net/addgene:26098; RRID:Addgene_26098)

## Data and materials availability

The authors confirm that the data supporting the findings of this study are available within the article and its supplementary materials.

## Supplementary Materials

**Supplementary Figure S1:**
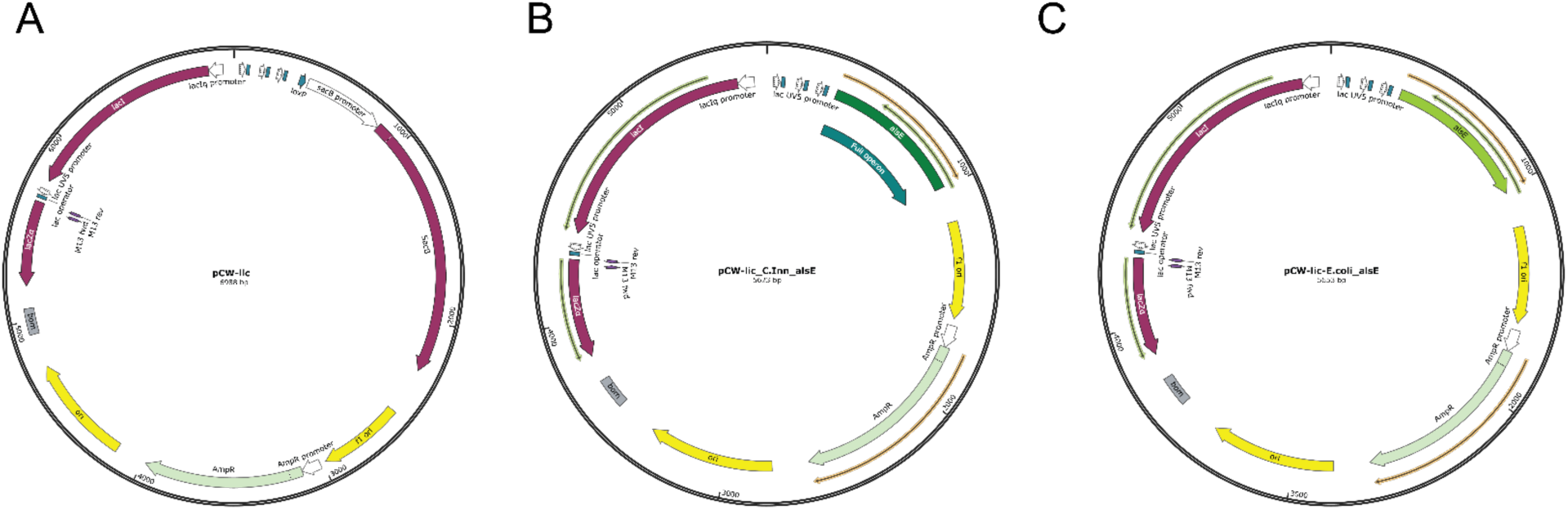
Plasmid maps of constructs. A) pCW-lic vector backbone.31 B) pCW-lic_C.inn_*alsE* vector containing the D-Allulose-6-phosphate 3-Epimerase (*alsE*) gene from *C. innocuum* ligated into the pCW-lic vector backbone via Gibson assembly. C) pCW-lic-E.Coli_*alsE* vector containing the *alsE* gene from *E. coli* ligated into the pCW-lic backbone. Pale green arrows represent an ampicillin resistance gene. Maps created using SnapGene.

**Supplementary Table 1:**
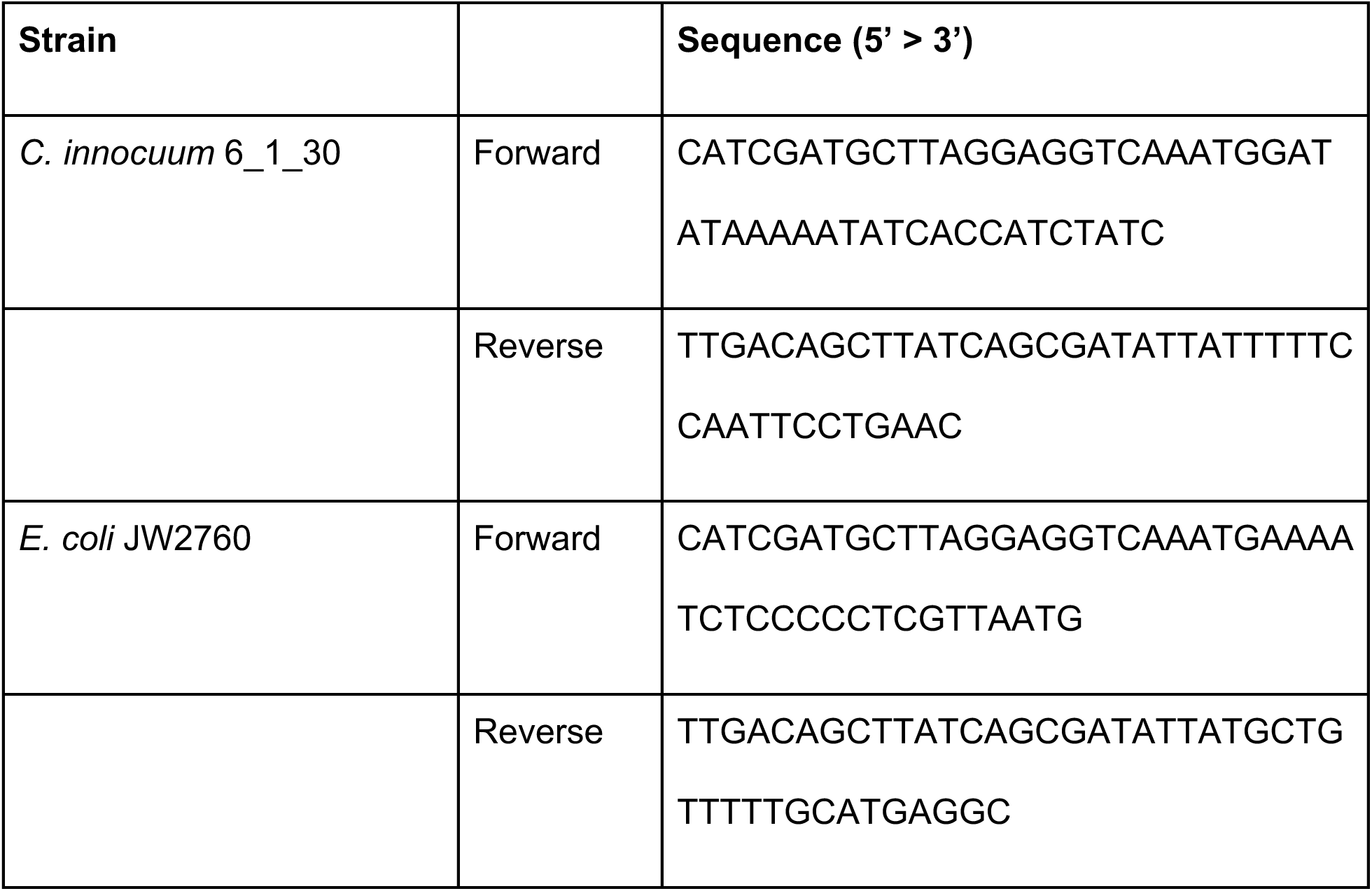
Primers used in construct development.

**Supplementary Table 2:** Predicted AlsE amino acid sequences identified with taxonomy information.

**Supplementary Table 3:** Metadata on the metagenomics samples used in this study, including SRA numbers and sample ids, along with the read mapping counts and total reads.

